# Fbxo7 promotes Cdk6 activity to inhibit PFKP and glycolysis in T cells

**DOI:** 10.1101/2021.11.05.467417

**Authors:** Rebecca Harris, Ming Yang, Christina Schmidt, Sarbjit Singh, Amarnath Natarajan, Christian Frezza, Heike Laman

## Abstract

Deregulated Fbxo7 expression is associated with many pathologies, including anaemia, male sterility, cancer, and Parkinson’s disease, demonstrating its critical role in a variety of cell types. Although Fbxo7 is an F-box protein that recruits substrates for SCF-type E3 ubiquitin ligases, it also promotes the formation of cyclin D/Cdk6/p27 complexes in an E3-ligase independent fashion. We discovered PFKP, the major gatekeeper of glycolysis, in a screen for Fbxo7 substrates. PFKP has been previously shown to be a critical substrate of Cdk6 for the viability of T-ALL cells. We investigated the molecular relationships between Fbxo7, Cdk6 and PFKP, and the functional effect Fbxo7 has on T cell metabolism, viability, and activation. Fbxo7 promotes Cdk6-independent ubiquitination and Cdk6-dependent phosphorylation of PFKP. Importantly Fbxo7-deficient cells have reduced Cdk6 activity, and haematopoietic and lymphocytic cell lines show a significant dependency on Fbxo7. Compared to WT cells, CD4^+^ T cells with reduced Fbxo7 expression show increased glycolysis, despite lower cell viability and activation levels. Metabolomic studies of activated CD4^+^ T cells confirm increased glycolytic flux in Fbxo7-deficient cells, as well as altered nucleotide biosynthesis and arginine metabolism. We show Fbxo7 expression is glucose-responsive at the mRNA and protein level, and we propose Fbxo7 inhibits PFKP and glycolysis via its activation of Cdk6.

## Introduction

Fbxo7 (F-box protein only 7) is a clinically important protein implicated in a variety of pathologies, including anaemia, cancer and Parkinson’s disease (1). Like other F-box proteins, Fbxo7 functions as a receptor for <u>S</u>kp1-<u>C</u>ullin1-<u>F</u>-box protein (SCF)-type E3 ubiquitin ligases; however, it also acts as scaffold for other multimeric proteins, notably the G1-phase cell cycle regulators Cdk6 and p27 (1). Fbxo7 selectively promotes Cdk6, but not Cdk4, binding to its activators, the D-type cyclins, increasing kinase activity (2). D-type cyclins and Cdk4/6 are core components of the cell cycle regulatory machinery which have in common the capacity to phosphorylate and inactivate the G1 checkpoint proteins of the retinoblastoma-associated protein family. The G1 phase cyclins and Cdks are often over-expressed in multiple cancers, including hematopoietic, breast, and brain tumours, and thus they are well-validated targets for inhibition by small molecules, such as palbociclib, ribociclib, and abemaciclib, where their inhibition ostensibly reduces cell cycle entry (3-5). More recently, unique functions of Cdk6, distinct from Cdk4, in regulating specific transcription factors and cellular functions are being reported (6-10). This includes a pro-survival activity of cyclin D3/Cdk6 activity attributed to its phosphorylation and inhibition of glycolytic enzymes, PFKP and PKM2, in T acute lymphoblastic leukaemia (T-ALL) cells and other tumours with high expression of Cdk6 (11).

Our studies to identify the binding partners and substrates of Fbxo7 revealed an overlapping set of candidates that were identified as cyclin D3/Cdk6 substrates, including PFKP, the prominent PFK1 isoform expressed in T-ALL cells (12-14). We investigated the molecular relationships between these three proteins, and the effect Fbxo7 has on T cell metabolism, viability, and activation. We found SCF^Fbxo7^ ubiquitinates PFKP; however, an interaction between Fbxo7 and PFKP does not require Cdk6. In contrast, Cdk6 interaction with PFKP is dependent on Fbxo7, and Fbxo7-deficient T cells have reduced Cdk6 activity. Cancer cell lines derived from blood and lymphocytic malignancies, which have high expression of Cdk6, show a significant dependency on Fbxo7. Both naïve and activated CD4^+^ T cells isolated from Fbxo7-deficient mice show reduced cell viability and activation. Metabolomic studies of activated CD4^+^ T cells demonstrate that cells with reduced Fbxo7 have increased glycolytic flux compared to WT cells, alongside alterations in purine and pyrimidine biosynthesis and arginine metabolism. Our data show Fbxo7 is a dose-dependent, glucose-responsive gene. We propose Fbxo7 regulates metabolism at the entry of glucose into glycolysis by enabling Cdk6 phosphorylation of PFKP.

## Results

### PFKP is a substrate for ubiquitination by Fbxo7

To identify Fbxo7-interacting proteins, a yeast two-hybrid (Y2H) assay was performed using aa 334-522 of Fbxo7, which spans the F-box domain and the substrate-binding proline rich region, as bait. One candidate identified was aa 385-784 of 6-phosphofructokinase (PFKP). To validate this interaction in mammalian cells, endogenous Fbxo7 was immunoprecipitated from the T-ALL cell line, CCRF-CEM. Both Fbxo7 isoforms 1 and 2 of 70kDa and 60kDa, respectively, were isolated and immunoblot analyses for PFKP confirmed a specific interaction between the endogenous proteins (Fig. 1A).

**Figure 1.**
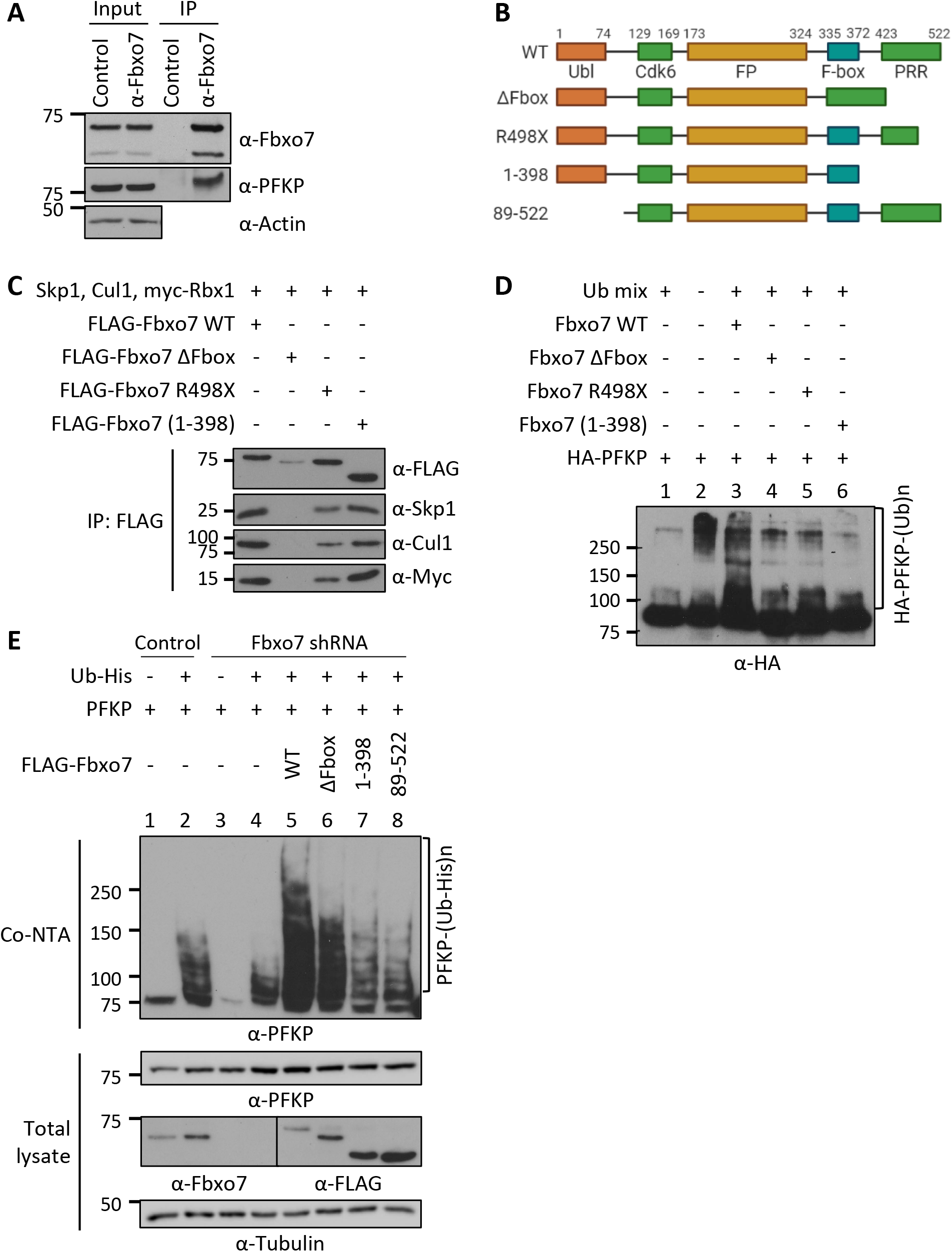
Fbxo7 ubiquitinates PFKP. **(A)** Fbxo7 immunoprecipitation from CCRF-CEM cells, showing co-immunoprecipitation of PFKP (n=3). **(B)** Schematic of Fbxo7 constructs used to make SCF ligases or for *in vivo* ubiquitination assays. All contain an N-terminal FLAG tag (not shown). Ubl, ubiquitin-like domain; Cdk6, Cdk6-binding domain; FP, Fbxo7-PI31 dimerization domain; PRR, proline rich region. **(C)** FLAG-Fbxo7 constructs were transfected into HEK293T cells alongside Skp1, Cullin1 (Cul1) and myc-Rbx1. SCF complexes were isolated by FLAG immunoprecipitation, and the presence of the other SCF components was confirmed by immunoblot. **(D)** *In vitro* ubiquitination assay of SCF^Fbxo7^ complexes in C, together with HA-PFKP and a ubiquitin mix (ubiquitin buffer, UBE1, UbcH5a and ATP) (n=2). **(E)** *In vivo* ubiquitination assay of Fbxo7 constructs with PFKP. Constructs were over-expressed in HEK293T cells stably expressing a control or Fbxo7 shRNA. Ubiquitinated proteins were isolated using a cobalt-NTA affinity resin (Co-NTA) to the His-tag on ubiquitin. Immunoblot for PFKP shows the degree of ubiquitination following expression of each Fbxo7 construct (n=3).

Since F-box proteins recruit substrates to SCF-type E3 ligases, we next investigated whether SCF^Fbxo7^ ubiquitinated PFKP. Various FLAG-Fbxo7 constructs were transfected into HEK293T cells along with additional SCF components to generate WT and truncated Fbox7 SCF E3 ubiquitin ligases, isolated by FLAG immunoprecipitation (Fig. 1B and C). These E3 ligases were used for *in vitro* ubiquitination assays with HA-purified PFKP as a substrate (Fig. 1D). Upon addition of SCF^Fbxo7^, we observed an increased smear of high MW bands representing poly-ubiquitinated PFKP (Fig. 1D). This modification was dependent on the F-box domain (Fig. 1D) which binds Skp1 and is required for SCF^Fbxo7^ complex formation (Fig. 1C). Although two C-terminally truncated Fbxo7 proteins recruited SCF-ligase components (Fig. 1C), these E3 ligases did not promote PFKP poly-ubiquitination (Fig. 1D), indicating the C-terminus of Fbxo7, as used in the Y2H screen, was required for PFKP ubiquitination.

We next performed *in vivo* ubiquitination assays in HEK293T cells expressing a control or Fbxo7-targeting shRNA. Cells were transfected with His-tagged ubiquitin, PFKP and FLAG-Fbxo7 constructs, and ubiquitinated proteins were isolated by cobalt-NTA affinity resin (Fig. 1E). Immunoblot analyses showed a reduction in His-ubiquitinated PFKP in cells expressing shRNA to Fbxo7 compared to control shRNA (Fig. 1E, lane 4 vs. lane 2). Re-expression of WT Fbxo7 markedly increased PFKP poly-ubiquitination (Fig. 1E, lane 5), and this was reduced when the ΔF-box, N- or C-terminal truncated Fbxo7 constructs were expressed (Fig. 1E, lanes 6-8). Together, these data show an interaction occurs between Fbxo7 and PFKP resulting in its ubiquitination by SCF^Fbxo7^, and this modification is dependent on both the N- and C-termini of Fbxo7.

### Fbxo7 promotes Cdk6-independent ubiquitination and Cdk6-dependent phosphorylation of PFKP

Cdk6 phosphorylates PFKP (11), and Fbxo7 both binds directly to and activates Cdk6 (2) and ubiquitinates PFKP (Fig 1D, 1E). Since phospho-degrons are a common recognition motif for SCF E3 ligases, we investigated whether Cdk6 or its kinase activity was required for Fbxo7 to interact with PFKP. This was tested by treating cells with two different Cdk6-specific PROTAC degraders (15, 16), or by treating cells with palbociclib. Inhibition of Cdk4/6 in palbociclib-treated cells was verified by immunoblotting cell lysates for phosphoSer780, a Cdk4/6 phospho-acceptor site in the retinoblastoma protein, and by profiling cell cycle parameters (Supplementary Fig. 1). CCRF-CEM cells were treated with vehicle control (DMSO), 1 μM palbociclib or 0.1 μM Cdk6 PROTAC for 24 hours, prior to Fbxo7 immunoprecipitation (Fig. 2A). PFKP co-immunoprecipitated with Fbxo7 in cells treated with palbociclib (Fig. 2A, lane 3) and in Cdk6-PROTAC-treated cells (Fig. 2A, lanes 4 & 5), demonstrating neither Cdk6 activity nor its presence was necessary for Fbxo7 binding to PFKP.

**Figure 2.**
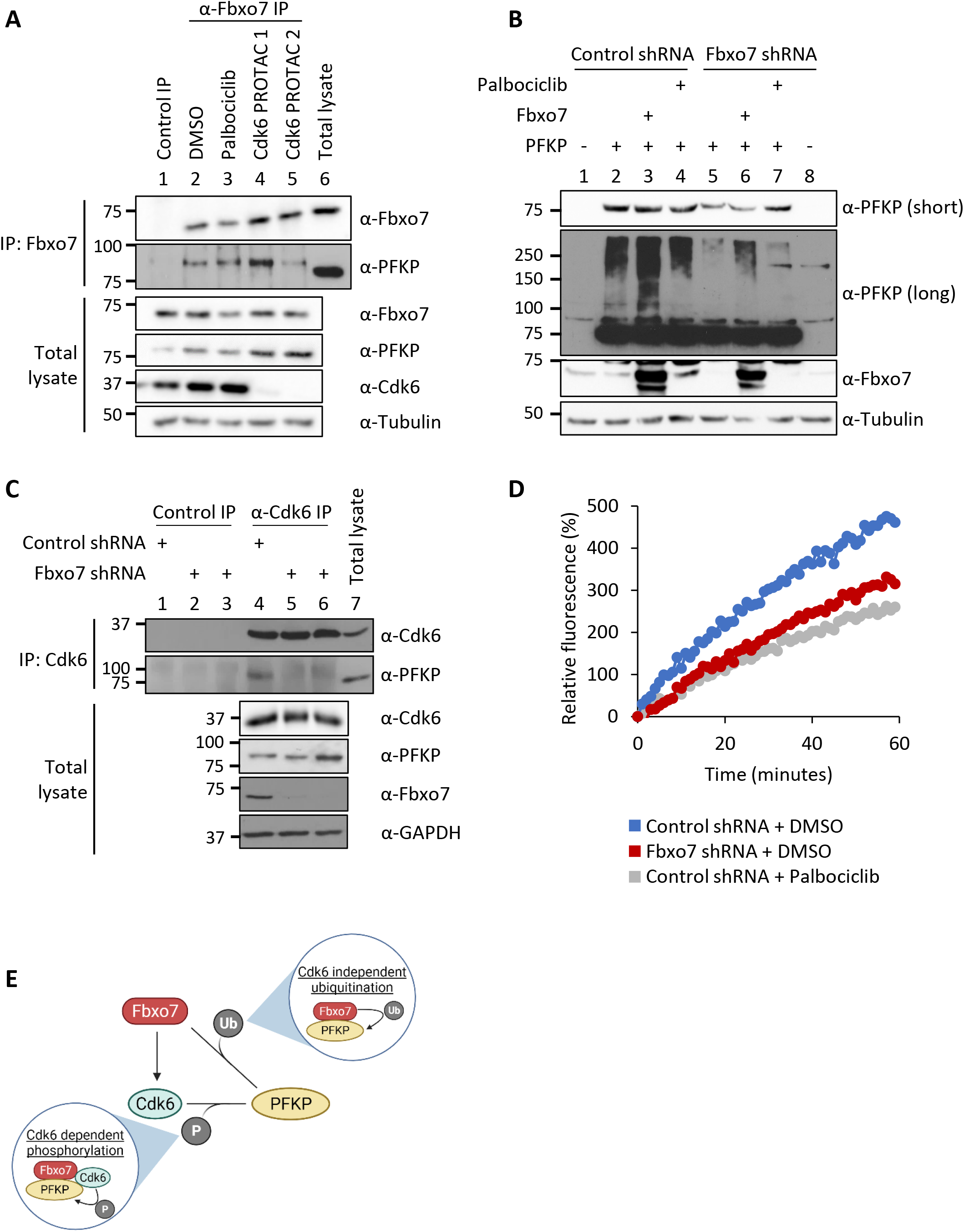
Fbxo7 promotes the Cdk6-independent ubiquitination and Cdk6-depedent phosphorylation of PFKP. **(A)** Fbxo7 immunoprecipitation from CCRF-CEM cells treated with 1 μM palbociclib or 0.1 μM of Cdk6 PROTAC for 24 hours. PFKP co-immunoprecipitation was analysed by immunoblot (n=3). **(B)** HEK293T cells with a control or Fbxo7 shRNA were transfected and treated with 1 μM palbociclib for 24 hours as indicated. Cell lysates were resolved by SDS-PAGE and analysed by immunoblot (n=2). **(C)** Cdk6 immunoprecipitation from CCRF-CEM cells expressing control or Fbxo7 shRNA. PFKP co-immunoprecipitation was analysed by immunoblot (n=1). **(D)** Cdk6 kinase activity in CCRF-CEM cells expressing control or Fbxo7 shRNA and treated with 1 μM palbociclib for 24 hours as indicated. Kinase activity is measured in a dose-dependent manner by fluorescence of the Cdk6 biosensor (n=1). **(E)** Diagram of model whereby Fbxo7 promotes both the ubiquitination and phosphorylation of PFKP.

We tested if Cdk6 activity was required for SCF^Fbxo7^-mediated ubiquitination of PFKP. HEK293T cells expressing a control or Fbxo7-targeting shRNA were transfected with PFKP and treated with palbociclib. As seen previously, PFKP ubiquitination was reduced upon Fbxo7 knockdown (Fig 2B, lane 2 vs. lane 5) and enhanced with Fbxo7 over-expression (Fig. 2B, lane 2 vs. lane 3). Moreover, 1 μM palbociclib treatment did not alter PFKP ubiquitination (Fig. 2B, lane 2 vs. lane 4), indicating Cdk6 activity is not required for SCF^Fbxo7^ ubiquitination of PFKP.

Since Cdk6 was dispensable for Fbxo7 interaction and ubiquitination of PFKP, we tested whether Fbxo7 was required Cdk6 interaction with PFKP. Cdk6 was immunoprecipitated from CCRF-CEM cells expressing control or two different Fbxo7-targeting shRNAs (Fig. 2C). Immunoblots show co-immunoprecipitation of PFKP with Cdk6 in the presence of Fbxo7 (Fig. 2C, lane 4), but this interaction is lost upon Fbxo7 knockdown (Fig. 2C, lanes 5 & 6). Similar results were obtained in MOLT-4 cells (Supplementary Fig. 2). These data show Fbxo7 is required for a Cdk6-PFKP interaction and suggest Fbxo7 would be required for Cdk6 phosphorylation of PFKP. To test this, we utilised a fluorescent peptide biosensor for Cdk6 activity based on the Cdk6-phosphoacceptor in PFKP (17). Lysates from WT or Fbxo7-deficient CCRF-CEM cells were assayed using this biosensor (Fig. 2D), and those with reduced Fbxo7 had lower Cdk6 activity, comparable to palbociclib-treated controls. These data show Fbxo7 is required for Cdk6 activity towards PFKP. Our data indicate Fbxo7 promotes two post-translational modifications, Cdk6 kinase-independent ubiquitination and Cdk6-dependent phosphorylation, on PFKP (Fig. 2E).

### Knockdown of Fbxo7 reduces the pool of inactive monomer/dimer forms of PFKP

One common outcome of ubiquitination is targeting to the 26S proteasome, so we tested whether Fbxo7 and PFKP protein levels correlated in CCRF-CEM and HEK293T cells. Immunoblot analyses of cell lysates showed no change in steady-state PFKP with either Fbxo7 knockdown or over-expression (Fig. 3A). There was also no change in PFKP half-life in cells with reduced Fbxo7 levels (Fig. 3B), and no accumulation of PFKP upon treatment with MG132 to inhibit the proteasome (Fig. 3C). These data indicate that Fbxo7 does not affect PFKP steady state levels. However, a major mechanism for PFKP regulation is the formation and dissociation of tetrameric complexes. PFKP is most enzymatically active as a tetramer, but not as dimers/monomers. We therefore investigated whether Fbxo7 expression affects the distribution of PFKP complexes. Total cell lysates from CCRF-CEM cells expressing a shRNA targeting Fbxo7 or a non-targeting control were passed over an ultrafiltration unit with molecular mass 200 kDa cutoff to separate monomers (86 kDa) and dimers (172 kDa) from active tetramers (344 kDa). As controls, allosteric regulators of PFKP were added to total cell lysates prior to filtration to promote (AMP) or dissociate (citrate) PFKP tetramers. The filtrates (<200kDa fraction) containing the smaller dimer/monomers forms were then analysed, alongside the lysates prior to filtration (Fig. 3D). Immunoblots showed that Fbxo7 knockdown did not change total levels of PFKP, as before, but instead reduced the amount of inactive dimer/monomer forms by 75% (Fig. 3D and 3E), suggesting a change in distribution of PFKP into active tetramers. We note that this phenotype is also observed when T-ALL cells were treated with palbociclib (11).

**Figure 3.**
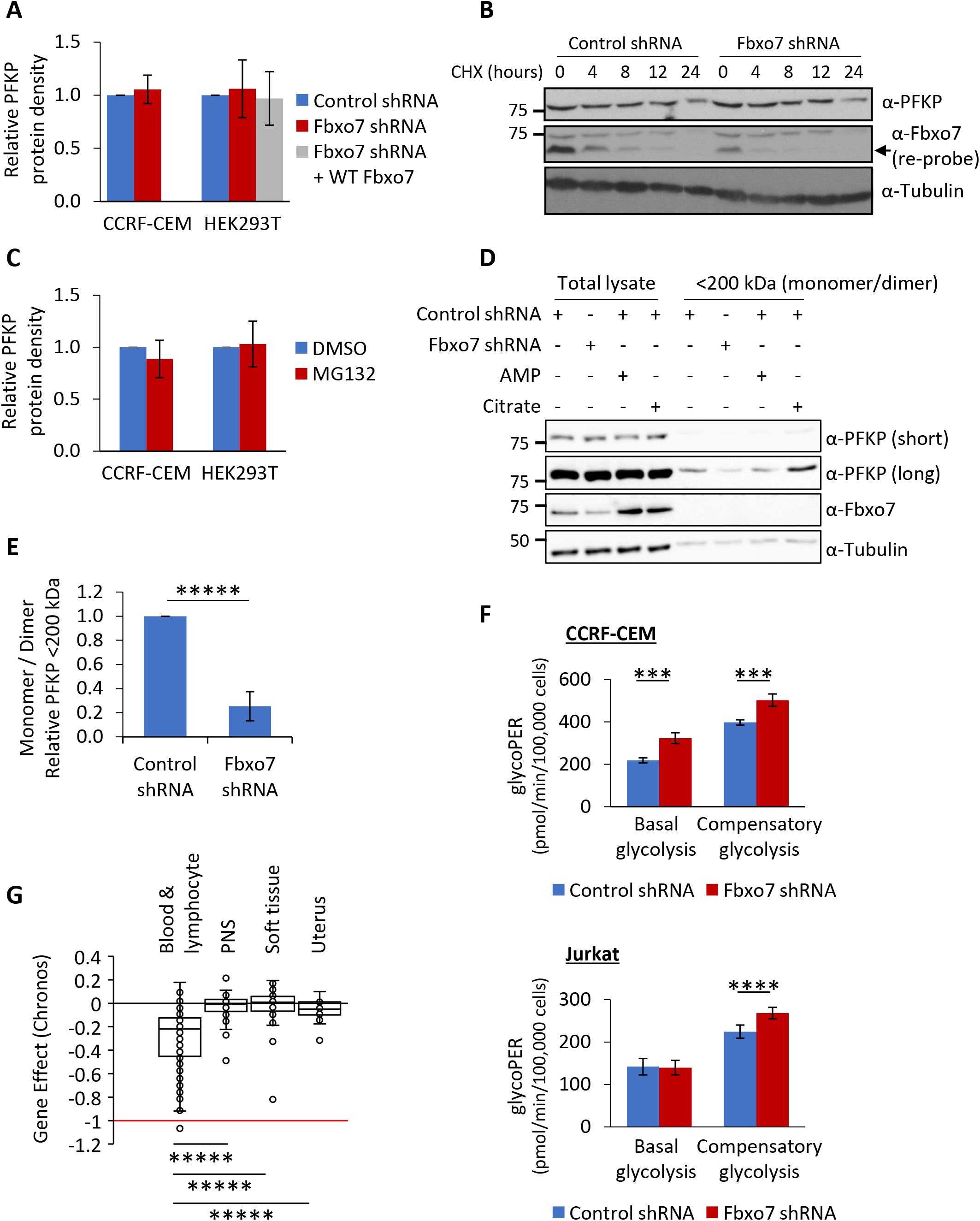
Fbxo7 knockdown in T-ALL cells reduces the proportion of inactive monomer/dimer PFKP and promotes glycolysis. **(A)** Lysates from cells expressing a control or Fbxo7 shRNA, and with Fbxo7 over-expression, were analysed by immunoblot and quantified (CCRF-CEM n=4, HEK293T n=3). **(B)** Representative immunoblot of PFKP half-life in CCRF-CEM cells expressing a control or Fbxo7 shRNA and treated with cycloheximide for up to 24 hours (n=2). **(C)** Cells were treated with DMSO or 10 μM MG132 for 4 hours and lysed. Proteins were resolved by SDS-PAGE to immunoblot for PFKP, and levels were quantified (CCRF-CEM n=2, HEK293T n=3). **(D)** Lysates from CCRF-CEM cells with control or Fbxo7 shRNA were passed over a 200 kDa molecular weight cut-off filter to separate PFKP monomers (86 kDa) and dimers (172 kDa) from tetramers (344 kDa). AMP and citrate were added to lysates as controls to respectively promote or dissociate PFKP tetramers. Total lysates and <200 kDa fractions were resolved by SDS-PAGE and analysed by immunoblot (representative n=4). **(E)** Quantification of PFKP in <200 kDa fraction as in D (n=4). **(F)** Glucose metabolism in CCRF-CEM (top) and Jurkat E6 (bottom) cells expressing a control or Fbxo7 shRNA was analysed by the Agilent Seahorse Glycolytic Rate Assay (n=4). **(G)** Chronos gene dependency scores for *FBXO7* in cancer cell lines of various lineage, plotted using data publically available on the DepMap portal (DepMap.org). A Chronos score of 0 indicates that a gene is non-essential, whilst -1 is comparable to the median of all pan-essential genes. (Blood & lymphocyte n=111, peripheral nervous system (PNS) n=32, soft tissue n=44, uterus n=34). *** p<0.005, **** p<0.001, ***** p<0.0005.

### Fbxo7 inhibits glycolysis in T-ALL cells and haematopoietic and lymphoid cancer cells are dependent on Fbxo7

As our data suggested that Fbxo7 increases PFKP activity, we tested whether Fbxo7 affected glycolysis in cancer cell lines using an Agilent Seahorse to measure the glycolytic rate. We measured the glycolytic rate of T-ALL cells expressing a control or Fbxo7 shRNA. We found Fbxo7 knockdown increased both basal and compensatory glycolysis in CCRF-CEM cells, by 48% and 26% respectively, whilst Jurkat E6 cells showed a 20% rise in compensatory glycolysis (Fig. 3F, Supplementary Fig. 3A). These data indicate higher levels of glycolysis in malignant T cells with reduced Fbxo7. Consequently, in light of studies showing Cdk6 regulation of glycolytic enzymes is essential for the survival of T-ALL cells (11), we reasoned that Fbxo7 should be essential for viability in haematological malignancies, since Cdk6 is the major kinase expressed in these cell lineages. We assessed the requirement for Fbxo7 in over 1000 cell lines by exploring the Cancer Dependency Map (18). Around 8% of cell lines were strongly selected as dependent on Fbxo7, and consistent with our model, we find that haematopoietic and lymphoid lineages are among the significantly enriched cell lines showing a preferential dependency on Fbxo7 (Fig. 3G, Supplementary Fig. 3B). These findings indicate an essential role for Fbxo7 in these malignancies.

### Metabolomic analysis on Fbxo7-deficient T cells shows broad metabolic alterations and increased glycolysis

To determine if Fbxo7 also increases glycolysis in primary T cells, where glycolysis is up-regulated upon activation, we tested the effect of Fbxo7 in T cells from WT or an Fbxo7-deficient mouse, in which the *Fbxo7* locus is disrupted by a *LacZ* insertion (19-21). WT and mutant splenic CD4^+^ T cells were isolated and activated *in vitro* for 48 hours prior to analysis (Fig. 4A, Supplementary Fig. 4A). Fbxo7-deficient T cells exhibited significantly higher basal glycolysis and a smaller increase in compensatory glycolysis (Fig. 4A, Supplementary Fig. 4A). In addition, we noted after activation, the viability of mutant T cells was significantly reduced by 1.9-fold compared to WT cells (Fig. 4B, Supplementary Fig. 4B and 4C), and the activation of Fbxo7-deficient cells was delayed and plateaued at 14.2% lower levels than WT (Fig. 4C). Thus, despite decreased viability and lower levels of activation, Fbxo7-deficient CD4^+^ T cells showed increased glycolysis compared to WT cells.

**Figure 4.**
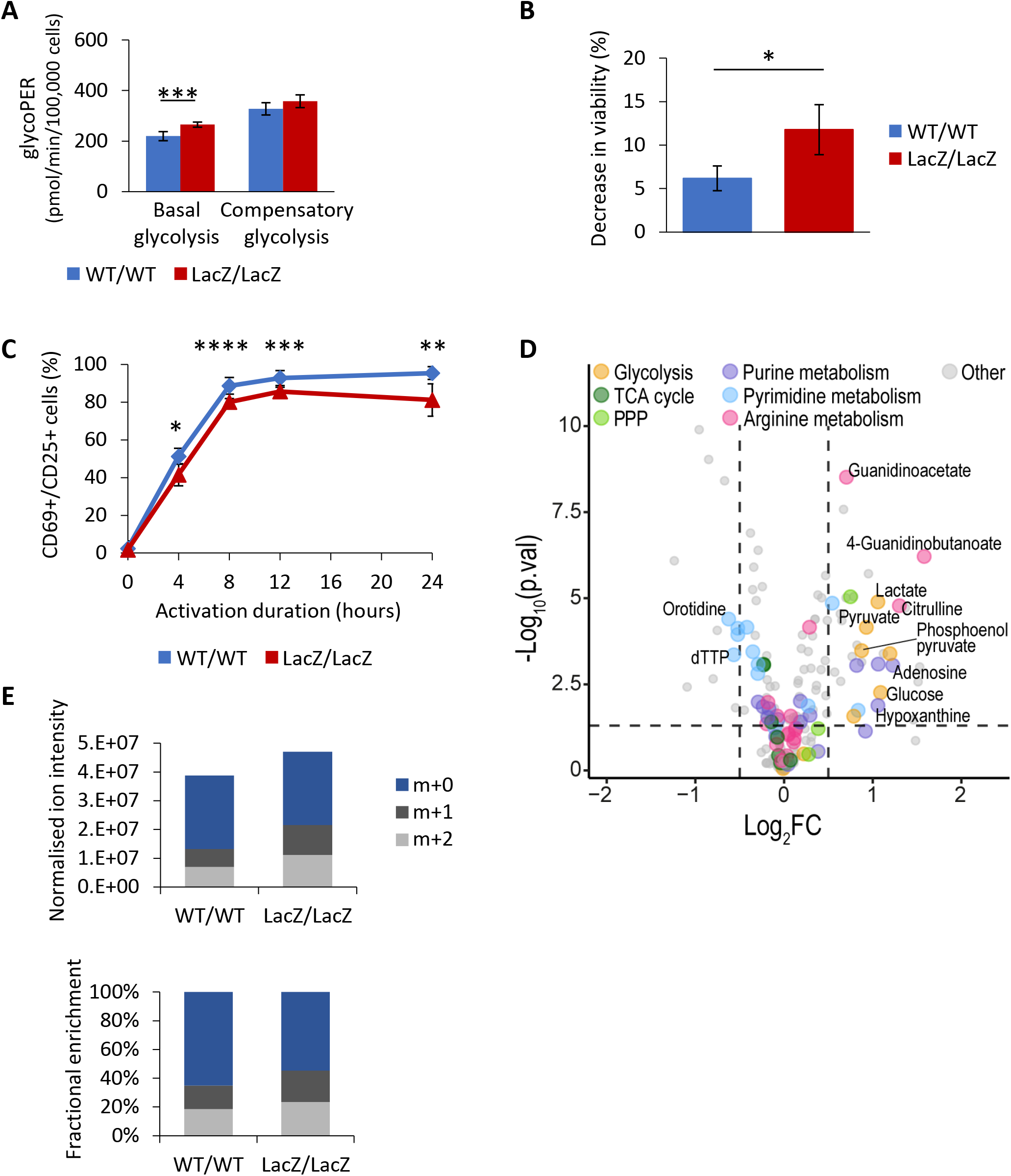
CD4^+^ T cells from mice lacking Fbxo7 display metabolic reprogramming and enhanced glycolysis, alongside survival and activation defects. **(A)** CD4^+^ T cells from WT or Fbxo7^LacZ/LacZ^ mice were activated for 48 hours, then their glucose metabolism was analysed by the Agilent Seahorse Glycolytic Rate Assay (n= 2). **(B)** Decrease in viability of CD4^+^ T cells over 24 hours of activation *in vitro* (n=3). **(C)** Activation of CD4^+^ T cells *in vitro*, as measured by CD69 and CD25 expression over 24 hours (n≥3). **(D)** Volcano plot of differentially expressed metabolites from untargeted metabolomics profiling of CD4^+^ T cells from WT or Fbxo7^LacZ/LacZ^ mice, activated for 48 hours (n=6). **(E)** CD4^+^ T cells from WT or Fbxo7^LacZ/LacZ^ mice were activated for 24 hours in the presence of glucose-1,2-^13^C_2_ then subjected to metabolomics analysis (n=6). m+1 and m+2 intracellular lactate are derived from radio-labelled glucose. * p<0.05, ** p<0.01, *** p<0.005, **** p<0.001.

To explore the impact of Fbxo7 on metabolism, we performed untargeted metabolomics profiling on activated splenic CD4^+^ T cells. Principal component analysis shows discrete clusters of WT and mutant samples, illustrating divergent metabolic profiles (Supplementary Fig. 4D). Variations in the T cells from mutant Fbxo7^LacZ/LacZ^ mice were multifaceted and included a dysregulation in arginine metabolism and a nucleotide imbalance caused by an increase in purines and reduction of pyrimidines (Fig. 4D). Importantly, there is a pronounced accumulation of lactate, pyruvate, and late-stage glycolytic intermediates in the Fbxo7-deficient T cells (Fig. 4D), supporting our hypothesis that glycolysis is altered in these cells.

To attribute this accumulation of glycolytic intermediates to increased glycolytic flux, we completed a stable isotope tracing study with CD4^+^ T cells activated for 24 hours in the presence of Glucose-1,2-^13^C_2_. There was a 63.4% increase in m+1 and m+2 labelled lactate in T cells from Fbxo7-deficient mice compared to WT (Fig. 4E), confirming an increase in lactate formation via glycolysis. These data reveal the broad metabolic alterations caused by a loss of Fbxo7 in CD4^+^ T cells and, crucially, highlight an increase in glycolytic flux, indicating Fbxo7 is a negative regulator of glycolysis in primary T cells. Overall, these data indicate higher levels of glycolysis in primary and malignant T cells with reduced Fbxo7, demonstrating Fbxo7 negatively regulates glycolysis.

### Glucose starvation induces Fbxo7 degradation by autophagy

The expression of glycolytic proteins is often responsive to glucose levels, so we tested whether Fbxo7 expression is influenced by glucose availability. HEK293T and three T-ALL cell lines were cultured in 4.5 g/L or 0 g/L glucose for 48 hours prior to lysis and immunoblotting. Fbxo7 protein was reduced by up to 65% following glucose starvation (Fig. 5A). Likewise, primary mouse CD4^+^ T cells activated for 48 hours in the absence of glucose showed a 67% decrease in Fbxo7 (Fig. 5A). Furthermore, there was a dose-dependent effect of glucose on Fbxo7 expression even up to hyperglycaemic levels (10g/L) (Fig. 5B). The reduction in Fbxo7 expression upon glucose withdrawal correlated with a 25% decrease in mRNA levels (Fig. 5C), and a shorter protein half-life as seen in a time course of CHX treatment of CCRF-CEM cells (*t*_*½*_ = 4 hr in 4.5g/L glucose vs. *t*_*½*_ = 1.75 hr in no glucose) (Fig. 5D). To determine the pathway promoting Fbxo7 degradation upon glucose removal, CCRF-CEM cells were treated with 10 μM MG132 or 200 nM bafilomycin A (BafA1) to inhibit the proteasome or autophagy, respectively (Fig. 5E). Surprisingly, proteasome inhibition with MG132 downregulated Fbxo7, but this was not dependent on glucose concentration. BafA1 treatment rescued Fbxo7 levels after glucose withdrawal, suggesting Fbxo7 was cleared by autophagy. These data show Fbxo7 expression correlates with glucose levels, and its regulation occurs both transcriptionally and post-translationally.

**Figure 5.**
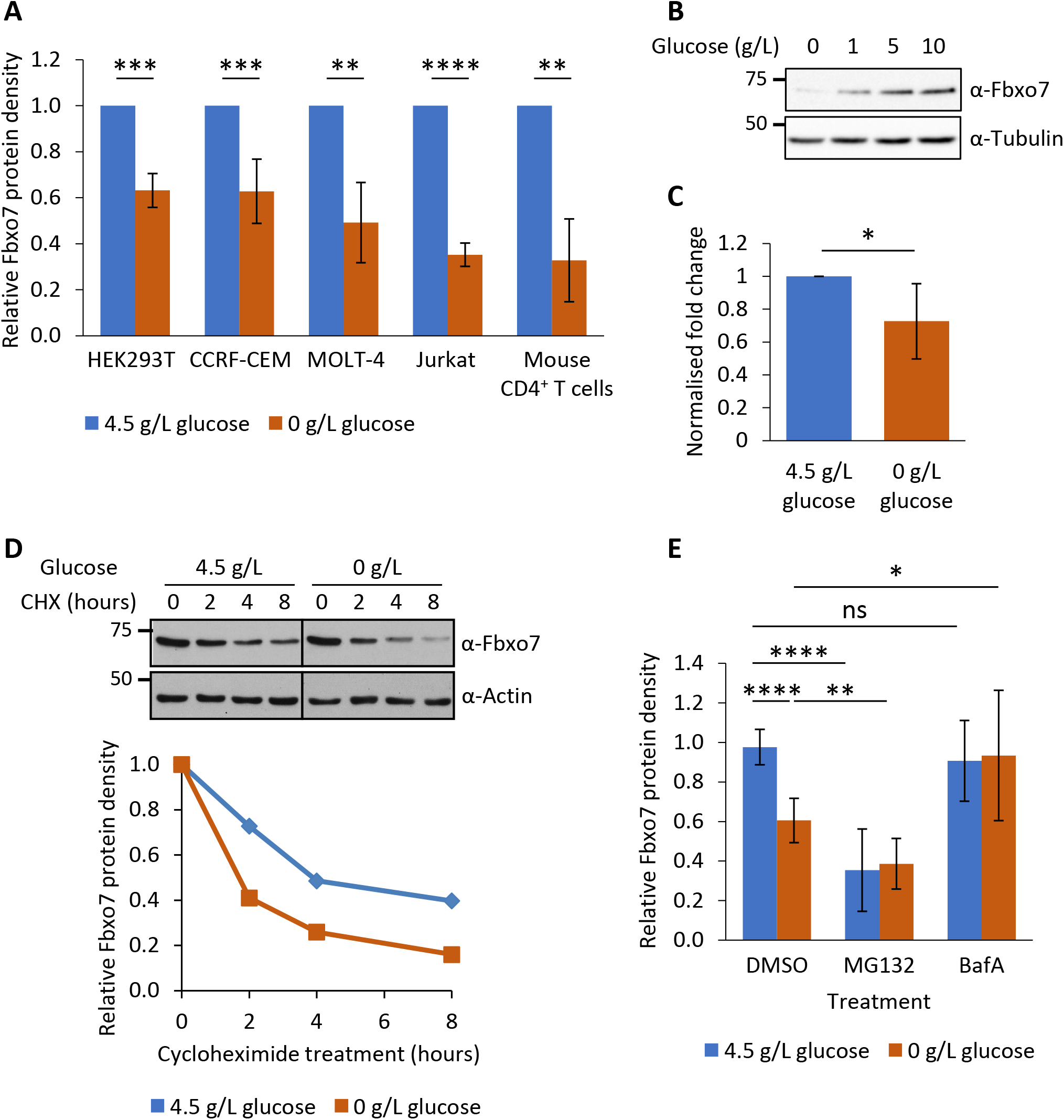
Regulation of Fbxo7 by glucose availability. **(A)** Relative Fbxo7 protein following 48 hours of glucose starvation, analysed by immunoblot and quantified (n≥3). **(B)** CCRF-CEM cells cultured for 48 hours with a titration of glucose, then lysed and analysed by immunoblot (n=3). **(C)** Relative Fbxo7 mRNA in naïve murine CD4^+^ T cells, incubated for 4 hours in 0 g/L or 4.5 g/L glucose (n=6). **(D)** Cycloheximide chase in CCRF-CEM cells following the removal (0 g/L) or maintenance (4.5 g/L) of glucose. Fbxo7 protein measured by immunoblot (top) and quantified (bottom) (representative of n=3). **(E)** CCRF-CEM cells were treated with DMSO, 10 μM MG132 or 200 nM BafA1 immediately following glucose removal (0 g/L) or maintenance (4.5 g/L). Fbxo7 protein measured by immunoblot and quantified (n=7). * p<0.05, ** p<0.01, ***p<0.001, **** p<0.0001

## Discussion

We show here Fbxo7 promotes two types of post-translational modifications on PFKP whereby Fbxo7 promotes Cdk6 kinase activity and can also independently ubiquitinate PFKP. We find several phenotypes observed when Fbxo7 is reduced in primary and malignant T cells, including decreases in the inactive dimer/monomer forms of PFKP, increases in glycolysis, and reduced T cell activation phenocopy the loss or inhibition of Cdk6 activity (11, 22-24). This supports the idea that a major role for Fbxo7 is Cdk6 activation, and our findings using a biosensor based on a Cdk6-phosphoacceptor in PFKP support this idea. Although PFKP was clearly ubiquitinated by SCF^Fbxo7^ independent of Cdk6 activity, reducing Fbxo7 did not alter the steady state levels of PFKP. However, other ubiquitin ligases, TRIM21 and A20, have been reported to catalyse ubiquitination-mediated degradation of PFKP in glioblastoma and PFKL in hepatocellular carcinoma (25, 26). Although the non-degradative effects of Fbxo7-mediated ubiquitination of PFKP in T cells remain to be elucidated, we hypothesize an essential role for Fbxo7 in cancer cell lines in blood lineages is as an activator of Cdk6. In support of this, we found that haematological and lymphocytic malignancies, where Cdk6 is highly expressed, are specifically dependent on Fbxo7.

Quiescent T cells predominantly use oxidative phosphorylation (OXPHOS) for their energy needs, while activated T cells increase glycolysis and glutaminolysis, alongside ROS production. Moreover, differentiated T cell subsets display discrete metabolic profiles, with effector T cells notably more glycolytic than regulatory and memory subsets. Interestingly, we show that activated Fbxo7-deficient CD4^+^ T cells have higher levels of glycolysis despite significantly lower levels of viability and activation, and an increase in the proportion of less glycolytic regulatory and central memory T cell subsets in mutant mice (27, 28). Our analysis of the metabolome of activated CD4^+^ T cells lacking Fbxo7 also indicates that, in addition to increases in glycolysis, these primary cells have numerous metabolic alterations, including in arginine metabolism and in purine and pyrimidine biosynthesis. The central role of arginine in T cell metabolism makes this an interesting area for future investigation (29).

Fbxo7 is a widely expressed protein, most highly in blood and testes. Fbxo7 mis-regulation and mutation are associated with a variety of pathological conditions, from anaemia, male sterility, cancer and Parkinson’s disease, demonstrating its critical role in multiple cell types. The diversity of these cell types raises the question of which Fbxo7-regulated pathways underlying cellular dysfunction are common or cell type-specific. We demonstrate that Fbxo7 expression is responsive to glucose levels at both the mRNA and protein levels, suggesting Fbxo7 levels may act to adjust the rate of glycolysis under different physiological and pathological conditions. We propose Cdk6 activity will respond to fluctuating glucose levels and potentially other stress conditions through the fine-tuning of Fbxo7 levels (30). In addition to regulating metabolism, another critical role for Cdk6 under stress conditions has been attributed to its regulation of transcription (31, 32). Bellutti, *et al*, proposed that Cdk6 protects cells from oncogenic transformation of bone marrow cells with BCR-ABL by suppressing p53-mediated stress responses, and in the absence of Cdk6, p53’s pro-apoptotic functions prevail (31). In a Parkinson’s disease mouse model with conditional loss of Fbxo7 in dopaminergic neurons, a p53 pro-apoptotic signature was detected (13). One possibility is the progressive neuronal death observed in this mouse arises from a lack of Fbxo7 and consequently Cdk6, to offset the p53 pro-apoptotic transcriptional response. Although Fbxo7 impacts on several other cellular pathways, as a factor that selectively scaffolds Cdk6, Fbxo7 levels may help to set a threshold for Cdk6-directed stress responses, transcriptional and metabolic, in many different cell types. Future efforts will be aimed at understanding whether the regulation of Cdk6 represents a common, therapeutically tractable, pathway mediating Fbxo7’s effects in physiological and pathological settings.

## Materials and Methods

### Resource availability

#### Lead contact

Further information and requests for resources and reagents should be directed to the lead contact, Dr Heike Laman.

#### Materials availability

- Plasmids generated in this study have been deposited to Addgene.
- The Fbxo7^LacZ^ mouse line (Fbxo7^tm1a(EUCOMM)Hmgu^) is available from the International Mouse Phenotyping Consortium.
- Cell lines are commercially available.

#### Data and code availability

Metabolomics data have been deposited at MetaboLights and are publicly available as of the date of publication. Accession numbers are listed in the key resources table. CRISPR–Cas9 screening results for DepMap version 21Q3 are publicly available at https://depmap.org (18). Any additional information required to reanalyse the data reported in this paper is available from the lead contact upon request.

### Experimental model details

#### Mice

All experiments in mice were performed in accordance with the UK Animals (Scientific Procedures) Act 1986 and ARRIVE guidelines. Animal licences were approved by the Home Office and the University of Cambridge’s Animal Welfare & Ethical Review Body Standing Committee. Experiments were performed under Home Office licence PPL 70/9001. Fbxo7 ^LacZ^ mice (Fbxo7 ^tm1a(EUCOMM)Hmgu^ on a C57BL/6J background) were bred as heterozygous crosses. WT, heterozygous and homozygous littermates were harvested between 6-9 weeks. Where littermates were not available, mice of a similar age were compared. Both male and female mice were used for experiments.

#### Cell lines

HEK923T cells and the human T-acute lymphoblastic leukaemia (T-ALL) cell lines CCRF-CEM, MOLT-4 and Jurkat E6, were purchased from ATCC. Cells were maintained in DMEM (HEK293T) or RPMI (T-ALL cells) supplemented with 10% heat-inactivated foetal bovine serum (Gibco), 100 U/mL penicillin and streptomycin (Gibco) at 37°C in a humidified 5% CO_2_ atmosphere. Cell lines stably expressing shRNA to human *FBXO7* were generated as described previously (33-35) and selected with 2 µg/mL puromycin (Sigma-Aldrich). To vary glucose concentration, cells were cultured in DMEM or RPMI without glucose, and glucose solution (Thermo Scientific) was supplemented as required.

### Method details

#### Antibodies

Antibodies against the following proteins were used for immunoblotting: Fbxo7 (generated in (2)), PFKP (CST 8164 or Abcam ab233109), Cdk6 (Santa Cruz sc-177), Skp1 (BD Biosciences 610530), Cullin1 (Santa Cruz sc-11384), Myc-tag (CST 2272), FLAG-tag (Sigma F3165), Actin (Sigma A2066), Tubulin (Sigma T6557), GAPDH (Sigma G9545), Phospho-Rb (Ser780) (CST 9307). Signal detection was by enhanced chemiluminescence (ECL) (GE Healthcare) or SuperSignal™ West Pico PLUS Chemiluminescent Substrate (Thermo Scientific).

#### DNA constructs

pSG5-HA-PFKP was kindly provided by Kyung-Sup Kim (Department of Biochemistry and Molecular Biology, Yonsei University) and subcloned into pcDNA3. pcDNA3 vectors expressing full length, truncated or ΔFbox FBXO7 with an N-terminal Flag tag have been previously described (36).

#### Immunoprecipitation

For immunoprecipitation, cells were lysed in Tween Lysis Buffer (50 mM HEPES, 150 mM NaCl, 0.1% Tween-20, 10% Glycerol, 1 mM EDTA, 2.5 mM EGTA, 1 mM DTT, 10 mM β-glycerophosphate) with a protease inhibitor cocktail (Sigma-Aldrich) and other inhibitors (1 mM PMSF, 10 mM NaF, 1 mM Na_3_VO_4_) and disrupted with mild sonication. Lysates were incubated with 0.4 μg anti-Cdk6 (Santa Cruz sc-177) or 1 μg anti-Fbxo7 (Aviva Systems Biology ARP43128_P050) antibodies, or isotype matched control, for 1 hour at 4°C with rotation, then 20 µL Protein A/G PLUS-Agarose (Santa Cruz) was added and samples were incubated for a further 2 hours. Beads were washed 4 times in lysis buffer and resuspended in 2XLaemmli loading buffer.

#### Purification of SCF^Fbxo7^ complexes and substrates

HEK293T cells were transfected with Skp1, Cullin1 and Myc-Rbx1, alongside FLAG-Fbxo7 constructs. After 48 hours, cells were resuspended in lysis buffer (50 mM Tris-HCl pH 7.5, 225 mM KCl, 1% NP-40) with a protease inhibitor cocktail (Sigma-Aldrich) and other inhibitors (1 mM PMSF, 10 mM NaF, 1 mM Na_3_VO_4_). Lysates were incubated with Anti-FLAG® M2 Affinity Gel (Sigma-Aldrich) for 4 hours at 4°C with rotation. Beads were washed 3 times in lysis buffer and twice in elution buffer (10 mM HEPES, 225 mM KCl, 1.5 mM MgCl_2_, 0.1% NP-40), and protein was eluted with 100 µg/mL FLAG peptide (Sigma-Aldrich) in elution buffer for 1 hour at 4°C with rotation. Purified SCF complexes were stored at -20°C in 15% glycerol.

#### *In vitro* ubiquitination assays

To purify substrate for *in vitro* ubiquitination, HEK293T cells were transfected with HA-PFKP which was immunoprecipitated with anti-HA agarose (Sigma-Aldrich) 48 hours after transfection. Substrate was eluted with 300 µg/mL HA peptide (Sigma-Aldrich) and stored at 20°C in 15% glycerol. For the *in vitro* ubiquitination assay, a ubiquitin mix was prepared with 100 nM E1 (UBE1, Bio-Techne), 500 nM E2 (UbcH5a, Bio-Techne), 20 µM human recombinant ubiquitin (Santa Cruz) and 2 mM ATP (Bio-Techne) in 1X ubiquitin conjugation reaction buffer (Bio-Techne). This was incubated for 5 min at RT then added to 100 nM SCF and 1 µL HA-PFKP substrate and incubated for 1 hour at 30°C. The entire 10 µL reaction was mixed with an equal volume of 2X Laemmli loading buffer, resolved by SDS-PAGE and analysed by immunoblotting.

#### *In vivo* ubiquitination assays

HEK293T cells were transfected with His-tagged ubiquitin, PFKP, and FLAG-Fbxo7 constructs, and cells were treated with 25 µM MG132 for 4 hours prior to lysis. Cells were harvested and a 10% portion was lysed in RIPA buffer (50 mM Tris pH 7.5, 150 mM NaCl, 1% NP-40, 0.5% sodium deoxycholate, 0.1% SDS) for the total lysate sample. The remaining cells were resuspended in CoNTA lysis buffer (6 M guanidinium-HCl, 100 mM Na_2_HPO_4_/NaH_2_PO_4_, 10 mM Tris-HCl pH 8, 5 mM imidazole, 10 mM β-mercaptoethanol) with 5 mM N-ethylmaleimide (Sigma), protease inhibitor cocktail (Sigma-Aldrich) and other inhibitors (1 mM PMSF, 10 mM NaF, 1 mM Na_3_VO_4_), and disrupted with mild sonication. Lysates were incubated with Super Cobalt NTA Agarose Affinity Resin (Generon) for 4 hours at 4°C with rotation. Beads were washed once in CoNTA lysis buffer, once in CoNTA wash buffer (8 M urea, 100 mM Na_2_HPO_4_/NaH_2_PO_4_, 10 mM Tris-HCl pH 6.8, 5 mM imidazole, 10 mM β-mercaptoethanol), and twice in CoNTA wash buffer containing 0.1% Triton-X 100, with an incubation of 5 min at RT between each wash. Beads were incubated in CoNTA elution buffer (200 mM imidazole, 150 mM Tris-HCl pH 6.8, 30% glycerol, 5% SDS, 720 mM β-mercaptoethanol) for 20 min at RT with shaking to elute his-tagged proteins. Both total lysate and eluate were resolved by SDS-PAGE and analysed by immunoblotting.

#### Drug treatments

Where indicated, cells were treated with 10 µM MG132 (Sigma), 200 nM bafilomycin A1 (Santa Cruz Biotechnology), 100 µg/mL cycloheximide (Sigma), 1 µM palbociclib (Sigma), or 0.1 µM Cdk6 PROTAC for the indicated durations. Cdk6 PROTAC 1 was kindly provided by Sarbjit Singh and Amarnath Natarajan (Eppley Institute for Cancer Research, University of Nebraska) (15), Cdk6 PROTAC 2 was kindly provided by Yu Rao and Zimo Yan (School of Pharmaceutical Sciences, Tsinghua University)(16).

#### Cdk6 activity assay

CCRF-CEM cells were collected and lysed in lysis buffer (PBS, 0.2% NP-40, 1 mM EDTA, protease inhibitor cocktail (Sigma-Aldrich) and 2 mM PMSF). Cells treated with 1 µM palbociclib (Sigma) for 24 hours were used as a control. 60 µg protein was plated into a 96 well Fluotrac™ 200 plate (Greiner) in triplicate. The assay was performed in 200 µL PBS supplemented with 5 mM MgCl_2_ and 0.5 mM ATP, and 200 nM Cdk6 peptide biosensor kindly provided by May Morris (Institut des Biomolécules Max Mousseron, University of Montpellier) (17). Changes in fluorescence emission of the TAMRA-labelled peptide biosensor were recorded at 30°C on a FLUOstar Omega (BMG) (excitation 560 nm / emission 590 nm). Biosensor fluorescence was subtracted from the samples containing protein lysate, and relative fluorescence was calculated.

#### 200 kDa cutoff ultrafiltration

5×10^7^ CCRF-CEM cells were collected per sample and lysed in lysis buffer (25 mM Tris-HCl pH 7.5, 150 mM NaCl, 1 mM EDTA, 5% glycerol, 1% NP-40, 10 mM β-glycerophosphate) with a protease inhibitor cocktail (Sigma-Aldrich) and other inhibitors (1 mM PMSF, 10 mM NaF, 1 mM Na_3_VO_4_). 1 mM AMP (Acros Organics) or 10 mM citrate (Sigma) were also added to the lysis buffer as controls where indicated. Equal amounts of protein were filtered using the Disposable Ultrafiltration Units with molecular weight 200 kDa cutoff (USY-20, Advantec MFS). 30 μg of total lysate or filtrate (<200 kDa) were resolved by SDS-PAGE and immunoblotted with the indicated antibodies.

#### Cancer Gene Dependency Scores

This study used DepMap 21Q3 dependency data for FBXO7, which had been generated from a CRISPR-Cas9 screen (available at https://depmap.org) (18). Samples were grouped by their predefined lineage, and lineages with fewer than 5 cell lines were excluded from analyses. Final analyses included 1020 cell lines.

#### Murine CD4^+^ T cell isolation & activation

Spleens were harvested from WT, heterozygous or homozygous Fbxo7^LacZ^ mice and processed to a single cell suspension. For n=2 of Fig. 3E, cells were incubated in RBC lysis buffer (eBioscience) for 5 minutes prior to centrifugation. For all other data, viable lymphocytes were separated with mouse Lympholyte® Cell Separation Media (Cedarlane Labs). CD4^+^ T cells were then isolated by negative selection using the MojoSort Mouse CD4 T Cell Isolation Kit (Biolegend). Isolated CD4^+^ T cells were seeded at 1×10^6^/mL in RPMI supplemented with 10% FBS, 100 U/mL penicillin and streptomycin and 5 µM β-mercaptoethanol. To activate, cells were added to plates coated with 2 µg/mL α-CD3 (clone 145-2C11) and containing 2 µg/mL soluble α-CD28 (clone 37.51) and incubated for the indicated duration. Cell viability and activation were measured by flow cytometry by staining with antibodies to CD4-PE (clone GK1.5), CD25-PE/Cy7 (clone PC61), CD69-FITC (clone H1.2F3) and an eFluor™ 780 fixable viability dye (eBioscience) for 30 min at 4°C in the dark. Samples were analysed on a CytoFLEX S flow cytometer.

#### Seahorse extracellular flux analysis

Cells were harvested and resuspended in Seahorse XF RPMI medium (Agilent), supplemented with 2 mM glutamine (Thermo Scientific), 10 mM glucose (Thermo Scientific) and 1 mM pyruvate (Thermo Scientific), at pH 7.4. Cells were plated into a Seahorse XF96 Cell Culture Microplate (Agilent) pre-coated with 22.4 µg/mL Cell-Tak solution (Corning) at 3×10^5^ cells/well and incubated at 37°C for 45-60 min in a non-CO_2_ incubator. Cells were analysed using the Seahorse XF Glycolytic Rate Assay Kit (Agilent) with of 0.5 µM rotenone/antimycin A (Rot/AA) and 50 mM 2-deoxy-D-glucose (2-DG). The assay was run at 37°C on a Seahorse XF96 analyser (Agilent) and data was analysed using Wave software (Agilent).

#### Metabolite extraction

For metabolite profiling (Fig. 4C & Supplementary Fig. 3C), murine CD4^+^ T cells were isolated and activated in a 12 well plate for 48 hours. For ^13^C stable isotope tracing (Fig. 4D), murine CD4^+^ T cells were isolated and activated in a 12 well plate with culture media containing 2g/L D-Glucose-1,2-^13^C_2_ (Sigma) for 24 hours. To extract metabolites, cells were harvested, washed twice in PBS and resuspended in 200 μL ice cold metabolite extraction solution (50% LC–MS grade methanol, 30% LC-MS grade acetonitrile, 20% ultrapure water, 5 μM valine-d8 as internal standard) per 1×10^6^ cells. Cells were incubated in a dry ice-methanol bath for 20 min, then at 4°C with shaking for 15 min. Samples were centrifuged at 13000 rpm for 20 min and the supernatant was collected into autosampler vials for LC-MS analysis.

#### Metabolite measurement by LC-MS

LC-MS chromatographic separation of metabolites was achieved using a Millipore Sequant ZIC-pHILIC analytical column (5 µm, 2.1 × 150 mm) equipped with a 2.1 × 20 mm guard column (both 5 mm particle size) with a binary solvent system. Solvent A was 20 mM ammonium carbonate, 0.05% ammonium hydroxide; Solvent B was acetonitrile. The column oven and autosampler tray were held at 40°C and 4°C, respectively. The chromatographic gradient was run at a flow rate of 0.200 mL/min as follows: 0–2 min: 80% B; 2-17 min: linear gradient from 80% B to 20% B; 17-17.1 min: linear gradient from 20% B to 80% B; 17.1-22.5 min: hold at 80% B. Samples were randomized and analysed with LC–MS in a blinded manner with an injection volume of 5 µl. Pooled samples were generated from an equal mixture of all individual samples and analysed interspersed at regular intervals within sample sequence as a quality control. Metabolites were measured with a Thermo Scientific Q Exactive Hybrid Quadrupole-Orbitrap Mass spectrometer (HRMS) coupled to a Dionex Ultimate 3000 UHPLC. The mass spectrometer was operated in full-scan, polarity-switching mode, with the spray voltage set to +4.5 kV/-3.5 kV, the heated capillary held at 320°C, and the auxiliary gas heater held at 280°C. The sheath gas flow was set to 25 units, the auxiliary gas flow was set to 15 units, and the sweep gas flow was set to 0 units. HRMS data acquisition was performed in a range of *m/z* = 70–900, with the resolution set at 70,000, the AGC target at 1×10^6^, and the maximum injection time (Max IT) at 120 ms. Metabolite identities were confirmed using two parameters: (1) precursor ion m/z was matched within 5 ppm of theoretical mass predicted by the chemical formula; (2) the retention time of metabolites was within 5% of the retention time of a purified standard run with the same chromatographic method.

#### Metabolomics data analysis

Chromatogram review and peak area integration were performed using the Thermo Fisher software Tracefinder (v.5.0). Correction for natural abundance was performed using the Accucor Package (v.0.2.3) (37) and the fractional enrichment was visualised using stacked bar graphs. For the total pools, the peak area for each detected metabolite was subjected to the “Filtering 80% Rule”, half minimum missing value imputation, and normalized against the total ion count (TIC) of the sample to correct any variations introduced from sample handling through instrument analysis. Sample were excluded after performing testing for outliers based on geometric distances of each point in the PCA score analysis as part of the muma package (v.1.4) (38). Afterwards, PCA analysis was performed using the R base package stats (v.4.0.5) (https://www.r-project.org/) with the function prcomp and visualised using the autoplot function of ggplot2 (v.3.3.3) (39) after loading the ggfortify package (v.0.4.11) (40, 41). Differential metabolomics analysis was performed using the R package “gtools”(v.3.8.2) (https://cran.r-project.org/web/packages/gtools/index.html) to calculate the Log2FC using the functions “foldchange” and “foldchange2logratio” (parameter base=2). The corresponding p-value was calculated using the R base package stats (v.4.0.5) with the function “t.test” (SIMPLIFY = F). Volcano plots were generated using the EnhancedVolcano package (v. 1.8.0) (42).

#### RT-qPCR

RNA extraction from murine CD4^+^ T cells was performed using the RNeasy Plus Mini Kit (Qiagen) and cDNA was generated using the QuantiTect Reverse Transcription Kit (Qiagen), both according to manufacturer’s instructions. RT-qPCR reactions were performed in triplicate using SYBR® Green JumpStart™ Taq ReadyMix™ (Sigma). Primers used were: Fbxo7 (forward CGCAGCCAAAGTGTACAAAG, reverse AGGTTCAGTACTTGCCGTGTG), Cyclophilin (forward CCTTGGGCCGCGTCTCCTT, reverse CACCCTGGCACATGAATCCTG), Actin (KiCqStart® mouse M1_Actb), HPRT (KiCqStart® mouse M1_Hprt). Reactions were run on an Eppendorf Mastercycler® ep realplex instrument and conditions were as follows: 95°C for 2 min; 45 cycles of 95°C for 15 sec, 58°C for 20 sec, 72°C for 15 sec, 76°C for 8 sec and read; followed by melting curve analysis from 65°C to 95°C to confirm product specific amplification. Relative gene expression was calculated using the Pfaffl analysis method with normalisation to three housekeeping genes (cyclophilin, actin, HPRT).

#### Cell cycle analysis

CCRF-CEM and MOLT-4 cells were incubated with 1 μM palbociclib for 24 hours. Cells were harvested, washed in PBS, and slowly resuspended in ice-cold 70% ethanol by vortexing. Cells were fixed for at least 24 hours, washed in PBS and stained with propidium iodide (PI). Samples were analysed on a Cytek DxP8 flow cytometer.

#### Quantification and statistical analysis

Immunoblot analyses were quantified using ImageJ processing software. Data are presented as mean ± standard deviation. Statistical differences were calculated using Student’s two-tailed t tests with a significant cutoff of p < 0.05.

#### Key resource table

Metabolomics repository accession number to be updated.

## Acknowledgements

We thank Klaus Okkenhaug and Anne Cooke for helpful discussions. We thank Yu Rao and Zimo Yan (School of Pharmaceutical Sciences, Tsinghua University) for providing the CP-10 Cdk6-PROTAC and May C. Morris (Institut des Biomolécules Max Mousseron, Montpellier, Languedoc-Roussillon, France) for providing the Cdk6 biosensor. This work is supported by funding from the CRUK Major Centre Cambridge Training Award.

## Author Contributions

All authors have read and approved the manuscript. RH carried out experiments, analysed data and wrote the manuscript. SS and AN provided Cdk6 PROTAC. MY, CS, and CF designed and analysed metabolomics experiments and data, and MY and CS performed metabolomics experiments. HL conceived of the study, analysed data, and wrote the manuscript.

## Figure Legends

**Supplementary Figure 1.**
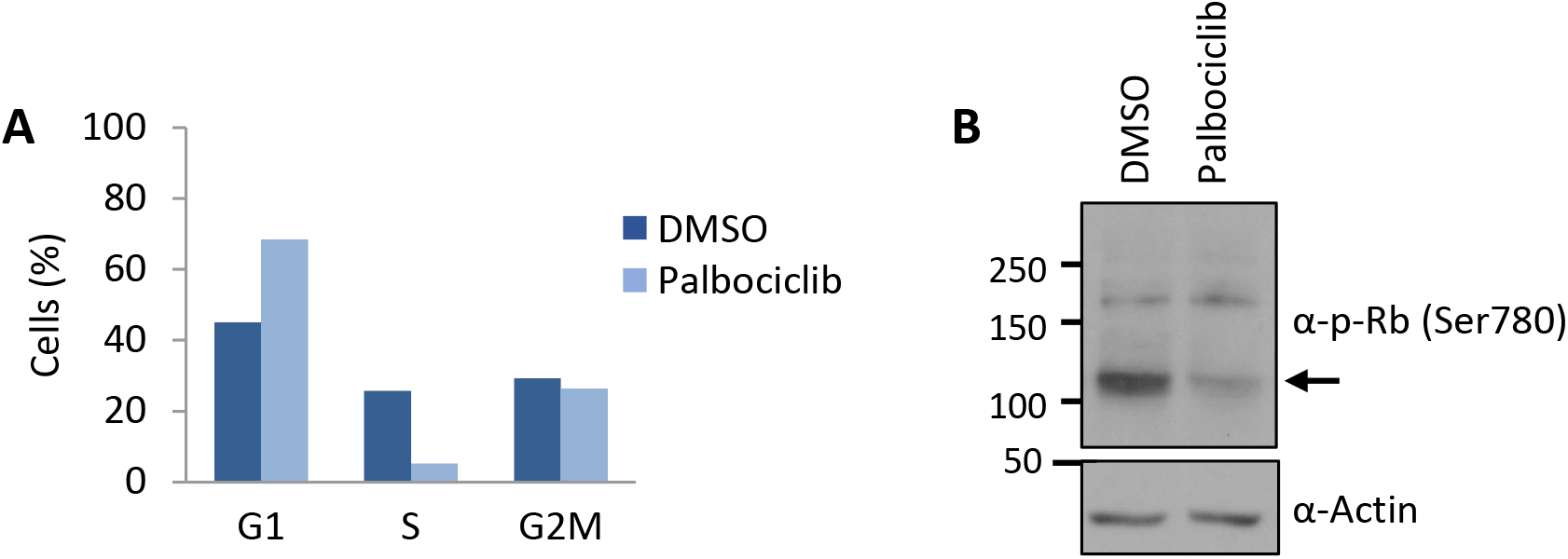
**(A)** Inhibition of Cdk4/6 by palbociclib in CCRF-CEM cells was verified by cell cycle analysis using propidium iodide staining (n=2). **(B)** Inhibition of Cdk4/6 by palbociclib in HEK293T cells was verified by immunoblotting cell lysates for phospho-Rb at Ser780 (n=1).

**Supplementary Figure 2.**
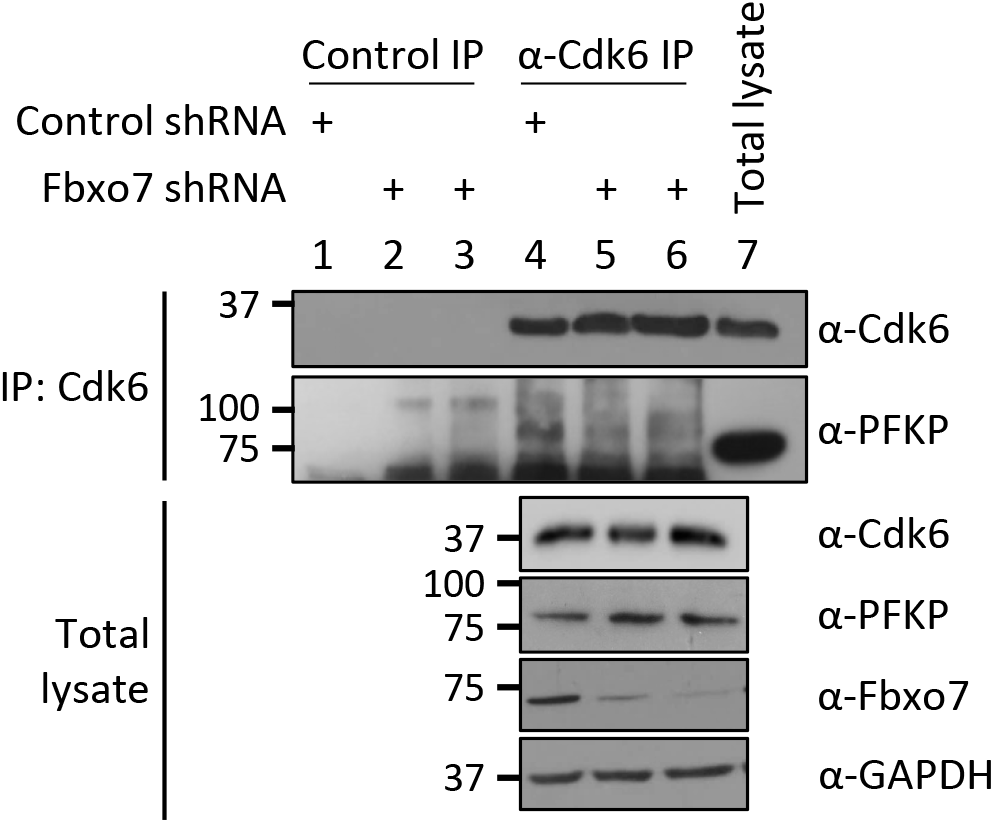
Cdk6 immunoprecipitation from MOLT-4 cells expressing control or Fbxo7 shRNA. PFKP co-immunoprecipitation was analysed by immunoblot (n=1).

**Supplementary Figure 3.**
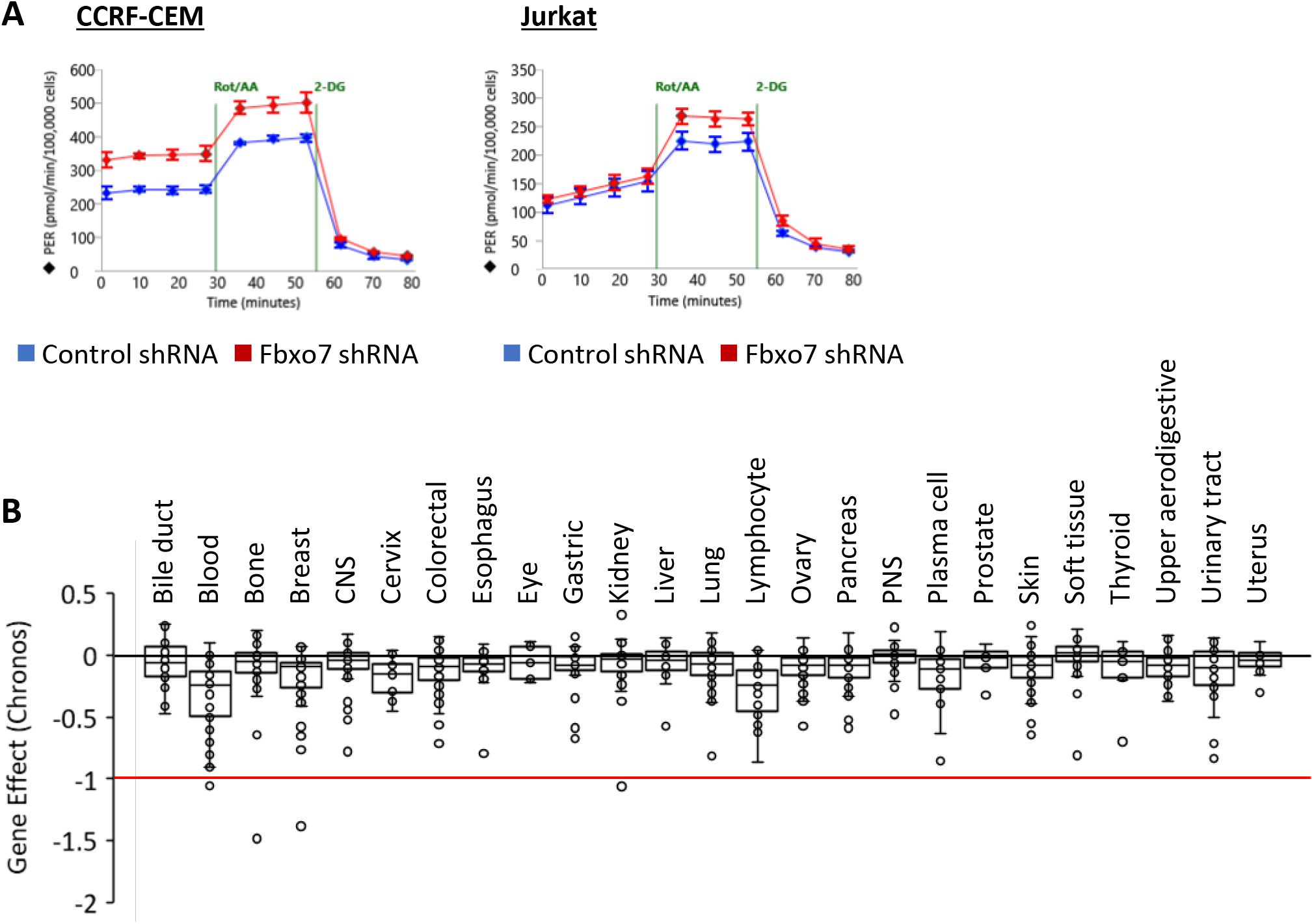
**(A)** Representative Seahorse profiles for CCRF-CEM (left) and Jurkat E6 (right) cells expressing a control or Fbxo7 shRNA and analysed by Agilent Seahorse Glycolytic Rate Assay. **(B)** Chronos gene dependency scores for *FBXO7* in 1020 cancer cell lines of various lineage, plotted using data publicly available on the DepMap portal (DepMap.org).

**Supplementary Figure 4.**
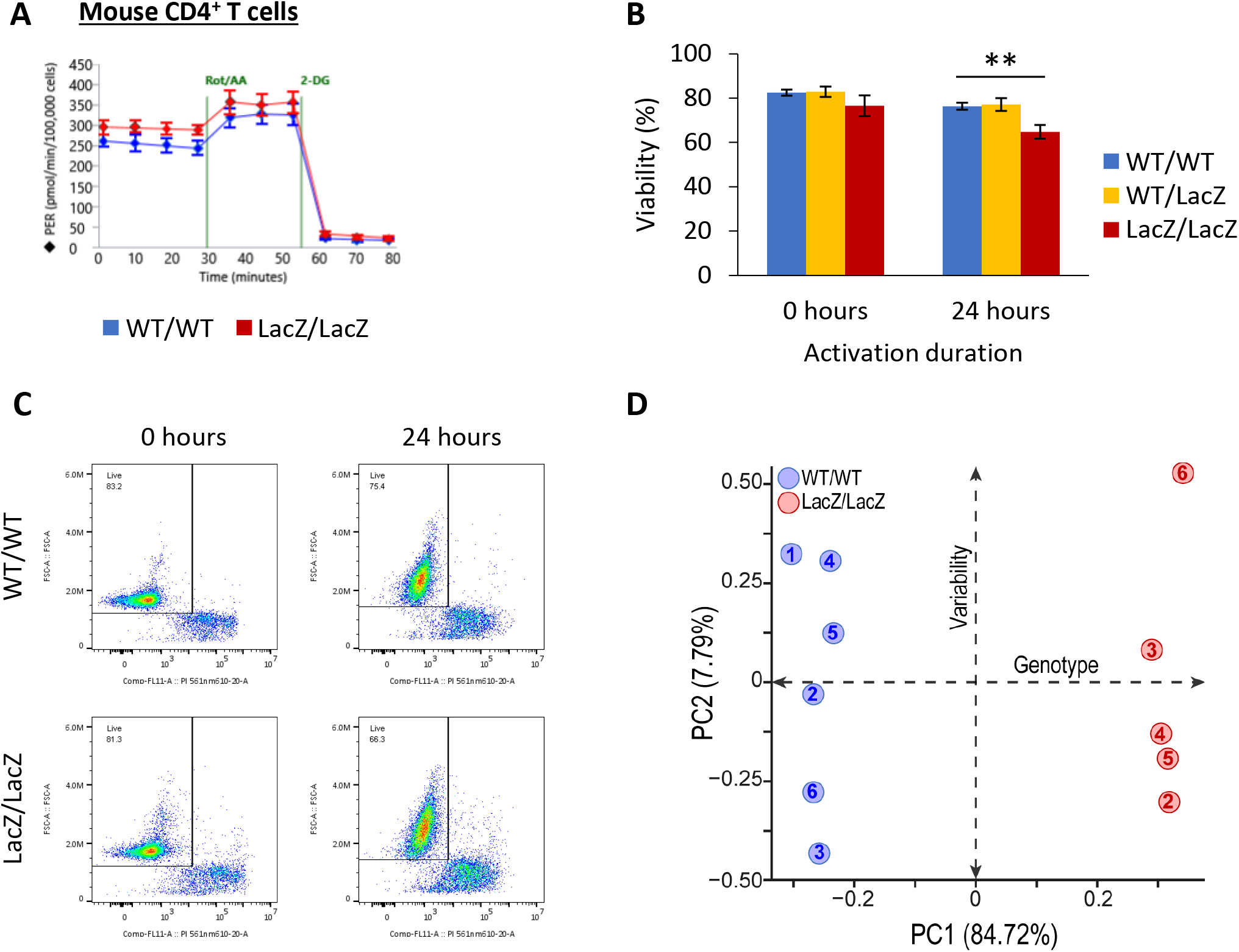
**(A)** Representative Seahorse profiles for CD4^+^ T cells from WT or Fbxo7^LacZ/LacZ^ mice activated for 48 hours and analysed by Agilent Seahorse Glycolytic Rate Assay. **(B)**- **(C)** Viability of CD4^+^ T cells upon isolation and 24 hours after activation *in vitro* (n=3). **(B)** Summary quantification. **(C)** Example FACS plots of PI staining. **(D)** Principal component analysis of intracellular metabolite ion intensities obtained from untargeted metabolomics profiling of CD4^+^ T cells from WT or Fbxo7^LacZ/LacZ^ mice, activated for 48 hours (n=6).

## References

1. Nelson DE, Randle SJ, Laman H. Beyond ubiquitination: the atypical functions of Fbxo7 and other F-box proteins. OpenBiol. 2013;3(10):130131.

2. Laman H, Funes JM, Ye H, Henderson S, Galinanes-Garcia L, Hara E, et al. Transforming activity of Fbxo7 is mediated specifically through regulation of cyclin D/cdk6. Embo J. 2005;24(17):3104–16.

3. Aleem E, Arceci RJ. Targeting cell cycle regulators in hematologic malignancies. Front Cell DevBiol. 2015;3:16.

4. Gao X, Leone GW, Wang H. Cyclin D-CDK4/6 functions in cancer. Adv Cancer Res. 2020;148:147–69.

5. Ingham M, Schwartz GK. Cell-Cycle Therapeutics Come of Age. J Clin Oncol. 2017;35(25):2949–59.

6. Uras IZ, Scheicher RM, Kollmann K, Glösmann M, Prchal-Murphy M, Tigan AS, et al. Cdk6 contributes to cytoskeletal stability in erythroid cells. Haematologica. 2017;102(6):995–1005.

7. Kollmann K, Heller G, Schneckenleithner C, Warsch W, Scheicher R, Ott RG, et al. A Kinase-Independent Function of CDK6 Links the Cell Cycle to Tumor Angiogenesis. Cancer Cell. 2016;30(2):359–60.

8. Scheicher R, Hoelbl-Kovacic A, Bellutti F, Tigan AS, Prchal-Murphy M, Heller G, et al. CDK6 as a key regulator of hematopoietic and leukemic stem cell activation. Blood. 2015;125(1):90–101.

9. Uras IZ, Walter GJ, Scheicher R, Bellutti F, Prchal-Murphy M, Tigan AS, et al. Palbociclib treatment of FLT3-ITD+ AML cells uncovers a kinase-dependent transcriptional regulation of FLT3 and PIM1 by CDK6. Blood. 2016;127(23):2890–902.

10. Tan C, Ginzberg MB, Webster R, Iyengar S, Liu S, Papadopoli D, et al. Cell size homeostasis is maintained by CDK4-dependent activation of p38 MAPK. Dev Cell. 2021;56(12):1756-69.e7.

11. Wang H, Nicolay BN, Chick JM, Gao X, Geng Y, Ren H, et al. The metabolic function of cyclin D3-CDK6 kinase in cancer cell survival. Nature. 2017;546(7658):426–30.

12. Teixeira FR, Randle SJ, Patel SP, Mevissen TE, Zenkeviciute G, Koide T, et al. Gsk3beta and Tomm20 are substrates of the SCFFbxo7/PARK15 ubiquitin ligase associated with Parkinson’s disease. BiochemJ. 2016.

13. Stott SR, Randle SJ, Al Rawi S, Rowicka PA, Harris R, Mason B, et al. Loss of FBXO7 results in a Parkinson’s-like dopaminergic degeneration via an RPL23-MDM2-TP53 pathway. JPathol. 2019.

14. Anders L, Ke N, Hydbring P, Choi YJ, Widlund HR, Chick JM, et al. A systematic screen for CDK4/6 substrates links FOXM1 phosphorylation to senescence suppression in cancer cells. Cancer Cell. 2011;20(5):620–34.

15. Rana S, Bendjennat M, Kour S, King HM, Kizhake S, Zahid M, et al. Selective degradation of CDK6 by a palbociclib based PROTAC. Bioorg Med Chem Lett. 2019;29(11):1375–9.

16. Su S, Yang Z, Gao H, Yang H, Zhu S, An Z, et al. Potent and Preferential Degradation of CDK6 via Proteolysis Targeting Chimera Degraders. J Med Chem. 2019;62(16):7575–82.

17. Soamalala J, Diot S, Pellerano M, Blanquart C, Galibert M, Jullian M, et al. Fluorescent Peptide Biosensor for Probing CDK6 Kinase Activity in Lung Cancer Cell Extracts. Chembiochem. 2021;22(6):1065–71.

18. Tsherniak A, Vazquez F, Montgomery PG, Weir BA, Kryukov G, Cowley GS, et al. Defining a Cancer Dependency Map. Cell. 2017;170(3):564-76.e16.

19. Ballesteros RC, Clare S, Arends MJ, Cambridge EL, Swiatkowska A, Caetano S, et al. FBXO7 sensitivity of phenotypic traits elucidated by a hypomorphic allele. PLoSOne. 2019;14(3):e0212481.

20. Randle SJ, Laman H. Structure and function of Fbxo7/PARK15 in Parkinson’s disease. CurrProtein PeptSci. 2016.

21. Meziane EKEK, Randle SJ, Nelson DE, Lomonosov M, Laman H. Knockdown of Fbxo7 reveals its regulatory role in proliferation and differentiation of haematopoietic precursor cells. Journal of Cell Science. 2011;124(2175-86):2175–86.

22. Hu MG, Deshpande A, Enos M, Mao D, Hinds EA, Hu GF, et al. A requirement for cyclin-dependent kinase 6 in thymocyte development and tumorigenesis. Cancer Res. 2009;69(3):810–8.

23. Malumbres M, Sotillo R, Santamaria D, Galan J, Cerezo A, Ortega S, et al. Mammalian Cells Cycle without the D-Type Cyclin-Dependent Kinases Cdk4 and Cdk6. Cell. 2004;118(4):493–504.

24. Hu MG, Deshpande A, Schlichting N, Hinds EA, Mao C, Dose M, et al. CDK6 kinase activity is required for thymocyte development. Blood. 2011;117(23):6120–31.

25. Lee JH, Liu R, Li J, Zhang C, Wang Y, Cai Q, et al. Stabilization of phosphofructokinase 1 platelet isoform by AKT promotes tumorigenesis. Nat Commun. 2017;8(1):949.

26. Feng Y, Zhang Y, Cai Y, Liu R, Lu M, Li T, et al. A20 targets PFKL and glycolysis to inhibit the progression of hepatocellular carcinoma. Cell Death Dis. 2020;11(2):89.

27. Patel SP, Randle SJ, Gibbs S, Cooke A, Laman H. Opposing effects on the cell cycle of T lymphocytes by Fbxo7 via Cdk6 and p27. Cell MolLife Sci. 2016.

28. Lomonosov M, Mezianee K, Ye H, Nelson DE, Randle SJ, Laman H. Expression of Fbxo7 in haematopoietic progenitor cells cooperates with p53 loss to promote lymphomagenesis. PLoSOne. 2011;6(6):e21165.

29. Geiger R, Rieckmann JC, Wolf T, Basso C, Feng Y, Fuhrer T, et al. L-Arginine Modulates T Cell Metabolism and Enhances Survival and Anti-tumor Activity. Cell. 2016;167(3):829-42.e13.

30. Zhou ZD, Sathiyamoorthy S, Angeles DC, Tan EK. Linking F-box protein 7 and parkin to neuronal degeneration in Parkinson’s disease (PD). Molecular Brain. 2016;9(1):41.

31. Bellutti F, Tigan AS, Nebenfuehr S, Dolezal M, Zojer M, Grausenburger R, et al. CDK6 Antagonizes p53-Induced Responses during Tumorigenesis. Cancer Discov. 2018;8(7):884–97.

32. Handschick K, Beuerlein K, Jurida L, Bartkuhn M, Müller H, Soelch J, et al. Cyclin-dependent kinase 6 is a chromatin-bound cofactor for NF-κB-dependent gene expression. Mol Cell. 2014;53(2):193–208.

33. Randle SJ, Nelson DE, Patel SP, Laman H. Defective erythropoiesis in a mouse model of reduced Fbxo7 expression due to decreased p27 expression. JPathol. 2015;237(2):263–72.

34. Burchell VS, Nelson DE, Sanchez-Martinez A, Delgado-Camprubi M, Ivatt RM, Pogson JH, et al. The Parkinson’s disease-linked proteins Fbxo7 and Parkin interact to mediate mitophagy. Nature neuroscience. 2013;16(9):1257–65.

35. Mezianee K, Randle SJ, Nelson DE, Lomonosov M, Laman H. Knockdown of Fbxo7 reveals its regulatory role in proliferation and differentiation of haematopoietic precursor cells. JCell Sci. 2011;124(Pt 13):2175–86.

36. Kirk R, Laman H, Knowles PP, Murray-Rust J, Lomonosov M, Mezianee K, et al. Structure of a conserved dimerization domain within the F-box protein Fbxo7 and the PI31 proteasome inhibitor. JBiolChem. 2008;283(32):22325–35.

37. Su X, Lu W, Rabinowitz JD. Metabolite Spectral Accuracy on Orbitraps. Anal Chem. 2017;89(11):5940–8.

38. Gaude E, Chignola F, Spiliotopoulos D, Spitaleri A, Michela Ghitti, Garcia-Manteiga JM, et al. muma, An R Package for Metabolomics Univariate and Multivariate Statistical Analysis. Current Metabolomics. 2013;1:180–9.

39. Hadley W. ggplot2: Elegant graphics for data analysis. : Springer-Verlag; 2016.

40. Horikoshi, M Y T. ggfortify: Data Visualization Tools for Statistical Analysis Results. 2018.

41. Tang, Y M H W L. ggfortify: Unified Interface to Visualize Statistical Result of Popular R Packages. The R Journal. 2016;8(2):474–85.

42. EnhancedVolcano: Publication-ready volcano plots with enhanced colouring and labeling. R package version 1.10.0, https://github.com/kevinblighe/EnhancedVolcano. 2021.

